# Phytoplankton Dynamics and Urbanization Gradients Reshape Dissolved Organic Matter and Microbial Networks across a Freshwater Estuary

**DOI:** 10.64898/2026.10.26.761939

**Authors:** Christian Ayala-Ortiz, Peter A. Birschbach, Josue Rodriguez-Ramos, William Petersen, Caroline Simonsen, Andrew Townsend, Izabel Stohel, Yuliya Farris, Aishwarya Santosh Deshpande, Anna Turetcaia, Thomas Wietsma, William Kew, Rosalie Chu, Euan D. Reavie, Christopher T. Filstrup, Hannah Nicklay, Emilly B. Graham

## Abstract

Freshwater estuaries process contaminants, cycle nutrients, and support biodiversity. However, watershed development degrades water quality, alters organic matter (OM) composition, and reshapes microbial and phytoplankton communities. Understanding how anthropogenic land use shapes estuarine ecosystem function remains a key challenge. We evaluated relationships among benthic and pelagic biogeochemistry, algal, and microbial communities across six sites along an urbanization gradient, combining FTICR-MS characterization of water and sediment OM, with 16S and 18S rRNA gene amplicon sequencing of bacterial/archaeal and algal assemblages, alongside measurements of nutrients, ions, turbidity, temperature, phytoplankton biovolume, and land use. Pelagic OM and microbial variation tracked seasonal gradients and transient phytoplankton dynamics modulated by urban inputs, whereas sediments responded directly to urbanization: urban sediments were N-rich with protein, and amino-sugar-like OM, while rural sediments carried aromatic, terrestrial, plant-derived signatures. Both compartments showed non-linear responses, with microbial, algal, and OM abundances shifting abruptly across urbanization thresholds. Strong associations between sediment microbial dissimilarity and phytoplankton biovolume indicate benthic–pelagic coupling, whereby urban nutrients putatively stimulate phytoplankton growth and subsequent OM deposition restructures benthic assemblages. Thus, pelagic communities capture transient urban effects mediated by phytoplankton, whereas benthic communities integrate chronic impacts, highlighting the value of compartment-specific monitoring in estuarine ecosystems.

## Introduction

Estuaries have outsized impacts on global biogeochemical processes due to the disproportionate amount of carbon and nutrients they contain. Internal estuarine biogeochemical dynamics depend largely on OM pools, which provide carbon and energy, and on the microorganisms that metabolize these pools. Human activities and rapid urban growth are degrading estuarine watersheds by altering nutrient delivery, organic matter (OM) concentration and composition, contaminant loads, and microbial and algal community dynamics (Lyu et al., 2021; Yang et al., 2017; Yunlin Zhang et al., 2016; Yutong Zhang et al., 2023; T. Zhao et al., 2025). However, the specific microbial taxa and organic molecules that are enriched or depleted as urbanization intensifies, and their impacts on estuarine biogeochemistry remain largely unknown. Because urban stressors produce complex responses, approaches integrating cross-domain interactions between aqueous geochemistry, microbial communities, and OM chemistry are required to understand their full effects on ecosystem function (X. Wang et al., 2024).

A key challenge in understanding the impacts of urbanization on estuaries is that land use may not affect all ecosystem compartments equally, as the water column and sediment are governed by different hydrobiogeochemical constraints (Hosen et al., 2017). As urbanization increases, hydrology and biogeochemical fluxes are altered by increased impervious surfaces, changed runoff timing, sediment loading, and enhanced nitrogen and phosphorus inputs (Barron et al., 2013; L. Yu et al., 2019; Yan Zhang et al., 2020). Estuaries are vulnerable to urbanization both directly through impervious-surface runoff that delivers nutrients, contaminants, and OM, and indirectly, by promoting nutrient enrichment and hydrologic modification that favor elevated phytoplankton biomass (L. A. Freeman et al., 2019). Water column impacts may be fleeting due to runoff events, storm surges, and wind-driven mixing, while sediments sequester longer term consequences of urbanization. This duality matters: if pelagic responses are episodic and reversible while benthic shifts are cumulative and potentially irreversible, then monitoring and management must be tailored to each compartment’s temporal dynamics. Identifying compartment-specific markers is therefore essential for monitoring and management (Hazen et al., 2024).

Urban runoff coincides with increased dissolved OM (DOM) concentration, changes in DOM composition, disrupted aquatic food webs, and altered global C and nutrient cycles (Xenopoulos et al., 2021). Because DOM composition, abundance, and bioavailability modulate the biogeochemical cycles in estuaries (Zhuang & Yang, 2018), land use-driven changes in the OM pool can fundamentally alter the biogeochemical context for microbial communities and the resultant microbially-mediated greenhouse gas emissions.

Furthermore, urbanization-driven nutrient inputs interact with seasonal warming and precipitation events to intensify phytoplankton responses (Birschbach et al., 2026). In urban estuaries, increases in phytoplankton biovolume rapidly transforms DOM pools through exudation, cell lysis, and microbial recycling (Eigemann et al., 2023; Luria et al., 2017). In turn, phytoplankton-derived OM can disrupt microbial communities and biogeochemical cycling (Chen et al., 2021; J. Zhao et al., 2021), and may settle into sediments, resulting in a benthic–pelagic coupling (Xiao et al., 2017). These responses appear to be non-linear, triggering ecological thresholds that reorganize microbial communities and DOM pools (Batt et al., 2013; Scheffer et al., 2009). Because phytoplankton abundance changes quickly over time and space (Ortiz et al., 2020), single measurements are insufficient. Connecting phytoplankton biovolume with shifts in algal communities, aquatic microbiomes, and DOM chemistry provides a clearer way to understand the causes and consequences of blooms.

Here, we examine how urbanization shapes estuarine biogeochemistry and algal, bacterial, and archaeal communities along a land-use gradient in the St. Louis River Estuary (SLRE), focusing on how phytoplankton mediates potential cross-domain interactions (i.e., bacteria/archaea - algae; bacteria/archaea - metabolite; or algae - metabolite) between water column and sediment habitats. We integrated Fourier-transform ion cyclotron resonance mass spectrometry (FTICR-MS) characterization of water-column and sediment OM, with 16S and 18S rRNA gene amplicon sequencing, alongside nutrient, ion, turbidity, temperature, and land-use data. We hypothesized that land use impacts estuarine ecosystems through both direct and indirect processes operating at fundamentally different temporal scales depending on the habitat (i.e., sediment vs. water column): fleeting seasonal forcings and phytoplankton variability would dominate water-column variability, whereas sediments would retain a stronger and more persistent urbanization imprint. We further hypothesized that urban land use and phytoplankton biovolume would drive distinct, directional shifts in OM composition towards XXXX; concomitant with restructuring of microbial communities. By combining habitat-specific analyses with indicator discovery approaches, this work (1) advances understanding of how urbanization and phytoplankton dynamics cascade through estuaries via compartment-specific processes, and (2) supports the development of actionable biomarkers for tracking phytoplankton-driven community reorganization and persistent sediment impacts in urban estuarine-freshwater networks.

## Methods

### Sample collection

We collected biweekly sediment and surface water samples at six different sites within the Lake Superior National Estuarine Research Reserve (LSNERR) from July to November 2023.

Surface water was collected using a Van Dorn water sampler at approximately 1.5 m depth below the surface. Approximately 1L was collected in a field-rinsed Nalgene bottle and used for subsampling, as follows. Filtered water was collected by attaching a 0.22 um Sterivex filter (MilliporeSigma, Burlington, MA, USA) to a 60-mL syringe. Approximately 5 mL of water was pushed through and discarded prior to sample collection to condition the filter. Two 15 mL conical tubes were filled with approximately 12 mL of filtered water for routine chemical analysis of cations and anions. A 40 mL amber vial was filled with filtered water by tilting the vial at approximately 45 degrees and gently dispensing water along the walls to minimize CO₂ degassing; minimal headspace (∼3 mm) was maintained for total organic carbon, inorganic carbon, and total nitrogen analysis. An additional 30 mL of filtered water was dispensed into a 60 mL amber vial pre-acidified with 85% phosphoric acid for molecular analysis.

Sediments were collected just below the waterline at approximately 0 - 10 cm depth using a clean, field-sterilized trowel. Each sediment was a composite from 3 to 5 different locations within a ∼ 2 m^2^ area. Sediments were collected into a 250-mL Nalgene and held at 4C between processing steps (below).

### Measurements of physicochemical properties

Water chemistry measurements were performed at the Environmental Molecular Sciences Laboratory in Richland, WA. Chlorophyll-a, conductivity, salinity, and turbidity were measured upon sample receipt using an AquaTROLL 500 multiparameter probe (in-Situ, Fort Collins, CO). pH was measured with a Mettler Toledo FiveEasy Plus pH sensor (Columbus, OH). The remaining water samples were stored at 4 °C until processing as follows. Anions (F^-^, Cl^-^, Br^-^) and cations (Li^+^, Na^+^, K^+^, Mg^+^, Ca^+^) were measured using a Thermo Scientific Dionex ICS-6000 ion chromatographer with a Dionex AS-AP refrigerated autosampler (Waltham, MA). Al, Ba, Cu, Fe, Mn, Mo, P, Si, Sr, and Zn were measured by an Agilent 8900 ICP-MS Triple Quad (Santa Clara, CA). A Shimadzu TOC-L/TN unit (Kyoto, Japan) was used to measure total organic and inorganic carbon (TOC and TIC) and total carbon and nitrogen (TC and TN) via catalytic combustion.

In sediments, gravimetric water content was measured in triplicate at EMSL by oven-drying ∼10g sediment at 100 °C for 24 hours and calculating the water loss. Extractable phosphorus was obtained using the Olsen or Bray method of extraction, depending on sediment pH. Where pH was ≥7.4, 3g of sediment was extracted with 20 mL of 0.5 M NaHCO3 solution adjusted to pH 8.5 with 2 M NaOH (Olsen). Where pH was <7.4, 2g of sediment was extracted with 14 mL solution of 0.025 N HCl and 0.03 N NH4F adjusted to pH 2.6 with 0.5 M HCl (Bray). Extracts were then shaken on a vortexer for 5 minutes at 1000 rpm, centrifuged at 4000 rpm for 20 minutes, and filtered through 0.45 μm Cytiva Whatman PES disk filters, pre-rinsed with MilliQ deionized water. The filtered extracts were stored at −20 °C until analysis. Phosphorus concentrations were measured colorimetrically using the Murphy-Riley ascorbic acid method (ammonium molybdate + antimony potassium tartrate + ascorbic acid), with absorbance measured at 882 nm either on an OI Analytical FS3700 with Teledyne Cetac autosampler or a non-automated Thermo GENESYS series spectrophotometer (Murphy and Riley, 1962) at Kuo Testing Laboratories in Pasco, WA.

To assay NO3-N, 8g of sediment was shaken with 24 mL of 0.5 M K2SO4 (1:3 w:v ratio) for 2 hours at 200 rpm, then centrifuged for 20 minutes at 4000 rpm. The supernatant was filtered, representing the salt-extractable, non-microbial fraction. This extract was stored at −20 °C until analysis for NO3-N (colorimetrically at 520 nm using a modified Cd-reduction method (Keeney and Nelson, 2015)) at Kuo Testing Laboratory in Pasco, WA.

Dissolved OM was obtained from sediments by shaking 6 g sediment with 30 mL MilliQ deionized water (1:5 w:v ratio) for 2 hours at 200 rpm, then centrifuging for 20 minutes at 4000 rpm. A portion of the supernatant was poured off for high resolution FTICR-MS analysis (see below), while the remaining was filtered using a 0.45 μm PES disk filter pre-rinsed with MilliQ deionized water, and then analyzed for water-extractable organic carbon and nitrogen. Measurements were done via catalytic combustion on a Shimadzu TOC-L Total Organic Carbon Analyzer with TNM-L Total Nitrogen Module. Water-extractable organic carbon was measured as non-purgeable organic carbon (NPOC).

Temperature and precipitation data were obtained from the Daymet V4 weather and meteorological summaries data collection (Thornton et al., 2020).

Any missing data was imputed using the Multivariate Imputation by Chained Equations (mice) approach using the mice R package (van Buuren & Groothuis-Oudshoorn, 2011).

### Phytoplankton enumeration and biovolume

Water samples for phytoplankton taxonomy were fixed and preserved with Lugol’s iodine for subsequent analysis following the quantitative Utermöhl settling chamber method (Utermöhl, 1958). For samples with high concentrations of protists or debris, a measured aliquot of sample water was placed in a Sedgewick-Rafter counting chamber. Counting of settled samples occurred on an inverted microscope (Olympus BX51) at 400× magnification. A minimum of 300 phytoplankton entities were counted per settled sample, and the transect area was recorded for quantitative calculations. For biovolume assessments, up to the first 10 specimens of each taxon observed were measured based on visible dimensions (length, width, diameter, depth, as necessary) and biovolume calculations for each taxon were conducted based on standardized cell shape formulae for each taxon (Reavie et al., 2010).

### Organic matter profiling via FTICR-MS

Organic matter profiling was completed using Fourier-transform ion cyclotron resonance mass spectrometry (FTICR-MS). Water samples and water extracts from sediments (see above) were diluted to a standardized 2 mg/L C and acidified to pH 2 using concentrated HCl (37 %). The extracts were then desalted by solid phase extraction(Dittmar et al., 2008) using a Gilson GX-274 liquid handler with Agilent Bond Elut PPL cartridges. Samples were eluted using 1.5 mL of Optima MeOH and transferred into 2 mL Microsolv vials. Post-SPE desalted samples were kept at −80 °C until analysis. Quality control checks were included with each batch of extractions, including SPE process blanks and standard Elliott Silt Loam soil (IHSS, Catalog # 1BS102M).

FTICR-MS spectra were acquired in negative mode with electrospray ionization (ESI) on a Bruker scimaX 7 Tesla FTICR mass spectrometer operating in quadrupolar detection mode, located at EMSL. Samples were infused at 4 μL/min using an automated flow injection system(Orton et al., 2018), with ESI high voltage of 3.3 kV, dry gas of 4 L/min at 200 °C. QC samples of MeOH (blanks) and Suwanee River Fulvic Acid (SRFA, obtained from IHSS) at 20 ppm in Optima MeOH were run at the start, end, and regularly throughout each sample set to monitor instrument performance. Spectra for each sample were acquired in triplicate. For each acquisition, ions were accumulated for 10 ms prior to transfer to the ICR cell, where spectra were acquired between m/z 107 and 1100. 300 transients (2.09s, 8MW) were co-added for each acquisition, yielding a typical magnitude mode resolving power at m/z 400 of ∼550k.

### FTICR-MS data processing and analysis

Raw FTICR-mass spectra were processed using CoreMS (Corilo et al., 2024), with a noise threshold of 10, and minimum peak prominence of 0.1. Mass calibration was performed to ensure each sample uses at least 200 calibration points. Molecular formula annotation was also performed with CoreMS, only the formula with the highest confidence score per each peak was kept. Peaks without assigned molecular formula were discarded. Data from replicate measurements was merged using molecular formula as the feature identity, only molecular formulas present in all available replicates were kept for further analysis. Molecular formulas were used to classify each peak into molecular classes based on H:C and O:C ratios using previous defined boundaries (Bailey et al., 2017; Tfaily et al., 2018).

### DNA extraction and amplicon sequencing

We extracted DNA from water and sediment samples using kit-based protocols with modifications to optimize yield.

For water samples, DNA was extracted from 0.22um Sterivex filters using the Qiagen DNeasy Blood and Tissue Kit following the manufacturer’s instructions (QIAGEN, Hilden, Germany). Whole filters were pushed to the bottom of the tube via pipette to ensure full submersion in the lysis solution. Following extraction, samples were eluted in 150 µL of elution buffer, which was passed through the filter membrane twice to maximize DNA recovery. All eluates were subsequently purified using the Zymo Clean and Concentrate Kit, including the 650 µL ethanol wash step, with a final elution passed through the column twice. Samples yielding less than 2 ng/µL were supplemented by extracting the corresponding duplicate filter using the initial elution as the final elution buffer to increase DNA concentration.

For sediment samples, DNA was extracted from 250 mg sediment using the FastDNA MPBio Kit (MP Biomedicals, Solon, OH, USA) with the following modifications: samples were homogenized using a bead beater at 30 m/s for 10 minutes, centrifugation in step 2 was extended to 15 minutes, 500 µL of supernatant was discarded in step 4, and a 15 mL tube was used in step 7. The Binding Matrix was vortexed prior to addition and fully resuspended when transferred from the tube to ensure complete DNA recovery. Following binding, 100 µL of elution buffer was added to the matrix and mixed thoroughly by repeated pipetting, then incubated at 55°C for 5 minutes on a heat block prior to the final elution. Eluates were purified using the Zymo Clean and Concentrate Kit as needed. Samples yielding less than 2 ng/µL were re-extracted and pooled to achieve sufficient concentrations for downstream analyses.

### Microbial community profiling via 16S and 18S rRNA gene sequencing

Amplicon library preparation and sequencing were performed by Azenta Life Sciences/GENEWIZ (South Plainfield, NJ, USA). Bacterial and archaeal 16S rRNA gene amplicons were generated using Azenta’s 16S-EZ primers and standard workflow, while eukaryotic 18S rRNA gene amplicons were generated using Earth Microbiome Project primers targeting the approximately 260-bp V9 region (±50 bp) (Amaral-Zettler et al., 2023). Samples were pooled at equimolar concentrations and sequenced via Illumina MiSeq using a 2 × 250-bp paired-end configuration, with a target depth of 20,000 amplicons per sample.

16S and 18S rRNA amplicons were processed using qiime2-amplicon version 2025.7 (Bolyen et al., 2019). Reads were imported and subsequently denoised using DADA2 (Callahan et al., 2016) with flags (16S) --p-trim-left-f 22, --p-trim-left-r 25, --p-trunc-len-f 0, --p-trunc-len-r 0, --p-n-threads 15, --p-trunc-q 2, and --p-chimera-method consensus and (18S) --p-trim-left-f 16, --p-trim-left-r 24, --p-trunc-len-f 170, --p-trunc-len-r 170, --p-trunc-q 2, --p-chimera-method consensus. Taxonomic assignment for both 16S and 18S rRNA amplicons was then performed with the classify-sklearn feature using the SILVA v138.2 NR99 uniform classifier from SILVA (Quast et al., 2013; Yilmaz et al., 2014). For the 18S amplicon dataset, detected ASVs were filtered to include only those from algal lineages.

### Statistical analysis and multi-omics integration

Analyses were performed separately for samples from the benthic and pelagic habitats. All statistical analyses were performed using R (v4.5.1) (R Core Team, 2025).

#### Alpha and beta diversity

For all datasets (FTICR-MS, 16S rRNA amplicon sequence variants (ASVs), and 18S rRNA algal ASVs) alpha diversity was estimated using the Chao1 richness estimator (Chao, 1984) and the Shannon diversity index (Shannon, 1948) using the vegan package (Oksanen et al., 2024). For the beta diversity analysis, Bray Curtis dissimilarity was calculated with the vegdist function (vegan package) using the normalized abundances of the features (i.e., molecular formulas or ASVs depending on the dataset) in each sample. To assess differences in DOM and microbial community composition across different sites and habitats, sample dissimilarities were visualized with a non-metric multidimensional scaling (NMDS) ordination using the vegan package. Measured biogeochemical parameters were fitted to the ordination using the envfit function with the purpose of identifying which environmental variables were significantly associated with the ordination axis aiding in the ecological interpretation of the ordination patterns. The relative contributions of urbanization (land-use gradient), temperature, and phytoplankton abundance to the compositional variation in each dataset (FTICR-MS, 16S, 18S) and compartment (water column, sediment) was assessed with a permutational multivariate analysis of variance (PERMANOVA) using the adonis2 function from vegan with 999 permutations.

#### Mantel tests

While envfit relates the environmental variables to the ordination axis, Mantel tests operate directly on the full dissimilarity matrices. Thus, they provide a complimentary insight by allowing to quantify the relationship between the environmental variables to the complete multivariate space resulting in a more robust support for the identified associations. Associations between specific environmental variables and the compositional dissimilarity of each sample were identified using Mantel tests correlating Bray-Curtis dissimilarity matrices from each dataset (FTICR-MS, 16S, and 18S) with Euclidean distances calculated from each individual measured environmental variable. The analysis was done using Pearson correlations and 999 permutations. Additionally, pairwise Pearson correlations between the environmental variables were computed and visualized with the ggcor package (Houyun Huang et al., 2020), and were used to contextualize the previous results.

#### Cross-domain association networks

The restructuring of ecological associations among bacteria/archaea, algae, and molecular formulas in the water column caused by the algal blooms was examined by constructing cross-domain association networks using the SparCC algorithm (Friedman & Alm, 2012) using the SpiecEasi package (Kurtz et al., 2025). For this, water-column samples were grouped into three categories based on phytoplankton biovolume: low (< 2.5 mm^3^/L), medium (2.5 - 5.5 mm^3^/L), and high (> 5.5 mm^3^/L). For each group, a separate SparCC network was constructed using the most prevalent (present in at least 3 samples for the 16S and 18S dataset, and in at least 12 samples for the FTICR-MS dataset) and abundant features of each dataset. Network edges were assessed using 100 bootstrapped estimates of the SparCC coefficients using the *sparccboot* function. Only edges with an absolute correlation coefficient in the upper 90th quantile, and whose sign was consistent across 80% of the bootstrapped estimates were retained in the final network.

Network properties including number of vertices, number of edges, average degree and connectivity were calculated using the *ggClusterNet* (Wen et al., 2025) package. Networks were visualized using the *igraph* (Csardi & Nepusz, 2005) and *ggnetwork* (Briatte, 2024) packages.

We restricted the cross-domain association network analysis to water-column samples because our central aim with this analysis was to resolve how algal blooms restructure the ecological associations among bacteria/archaea, algae, and DOM molecular formulas. As blooms are fundamentally a water-column phenomenon, and because we could directly stratify these samples along a measurable phytoplankton biovolume gradient (low, medium, and high), the water column provided the appropriate setting to capture bloom-driven shifts in cross-domain associations.

#### Threshold indicator taxa analysis (TITAN2)

Individual taxa and DOM molecules exhibiting consistent, directional responses along environmental gradients were identified using a Threshold Indicator Taxa Analysis (TITAN2) (Baker et al., 2025). This approach was applied separately to two gradients: (1) the phytoplankton biovolume gradient in water-column samples, to identify molecular and microbial indicators of bloom intensification; and (2) the urbanization gradient in sediment samples, to identify indicators of persistent land-use impacts. For each gradient, these analysis was applied independently to the FTICR-MS, 16S, and 18S datasets using the titan function from the TITAN2 R package (Baker et al., 2025) using 500 bootstrap replicates and only considering features that appeared in at least 5 samples.

#### Multiomics integration

Cross-domain feature sets capable of discriminating among bloom states or urbanization levels were identified using a multiblock sparse partial least squares discriminant analysis (Lê Cao et al., 2011) using the mixOmics package (Rohart et al., 2017). This supervised multivariate method integrates multiple data blocks (i.e., FTICR-MS, 16S, and 18S) to select a reduced set of discriminant features from each block that maximally separate predefined sample groups (i.e., phytoplankton groups and land-use groups for the water column and the sediment samples respectively).

We applied TITAN2 and multiblock sPLS-DA as complementary approaches to identify robust indicators of phytoplankton biovolume: TITAN2 was used to detect taxa and DOM molecules that respond consistently along the phytoplankton gradient and to pinpoint the specific thresholds at which these community reorganizations occur, while multiblock sPLS-DA was used to determine whether a compact subset of discriminant features could classify samples according to their phytoplankton biovolume. Using both methods allowed us to assess the concordance between threshold-based and discriminant-based indicators, thereby strengthening confidence in the features identified

## Results

### Habitat-specific processes shape variation in dissolved organic matter and microbial community composition

To evaluate how urbanization shapes benthic and pelagic DOM and microbial composition, we combined DOM characterization (FTICR-MS), with bacterial/archaeal (16S rRNA) and algal (18S rRNA) amplicon sequencing. Then, we applied multivariate statistics to assess which biogeochemical factors explained the observed compositional variation.

We detected 43,555 DOM molecules across all samples (n = 64), with water samples containing more molecules on average than sediments (Figure 2B). Both Chao1 richness and Shannon diversity indicated higher DOM pool diversity in the water column than in sediments (Figure 2A). Putative molecular class assignment based on H:C and O:C ratios showed that water DOM was dominated by lignin-like compounds, while sediments were dominated by a mixture of lignin-like and protein-like compounds (Figure 3A).

**Figure 1.**
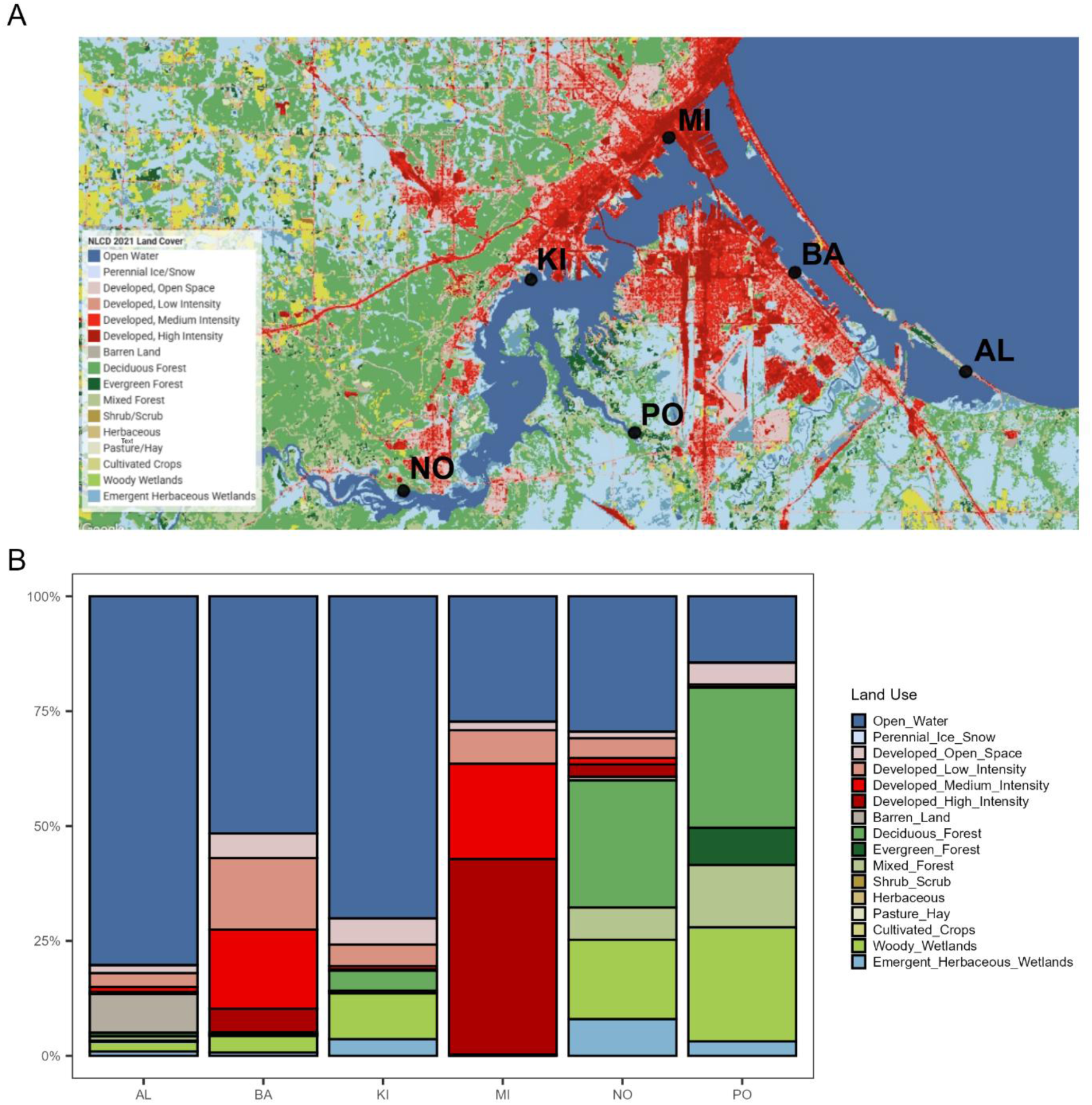
A) Map of the six analyzed sites showing the land use coverage based on the USGS National Land Cover Database, 2021 release. B) Bar plot showing the percentage of land use coverage in a radius of 500 m around the sampling sites.

**Figure 2.**
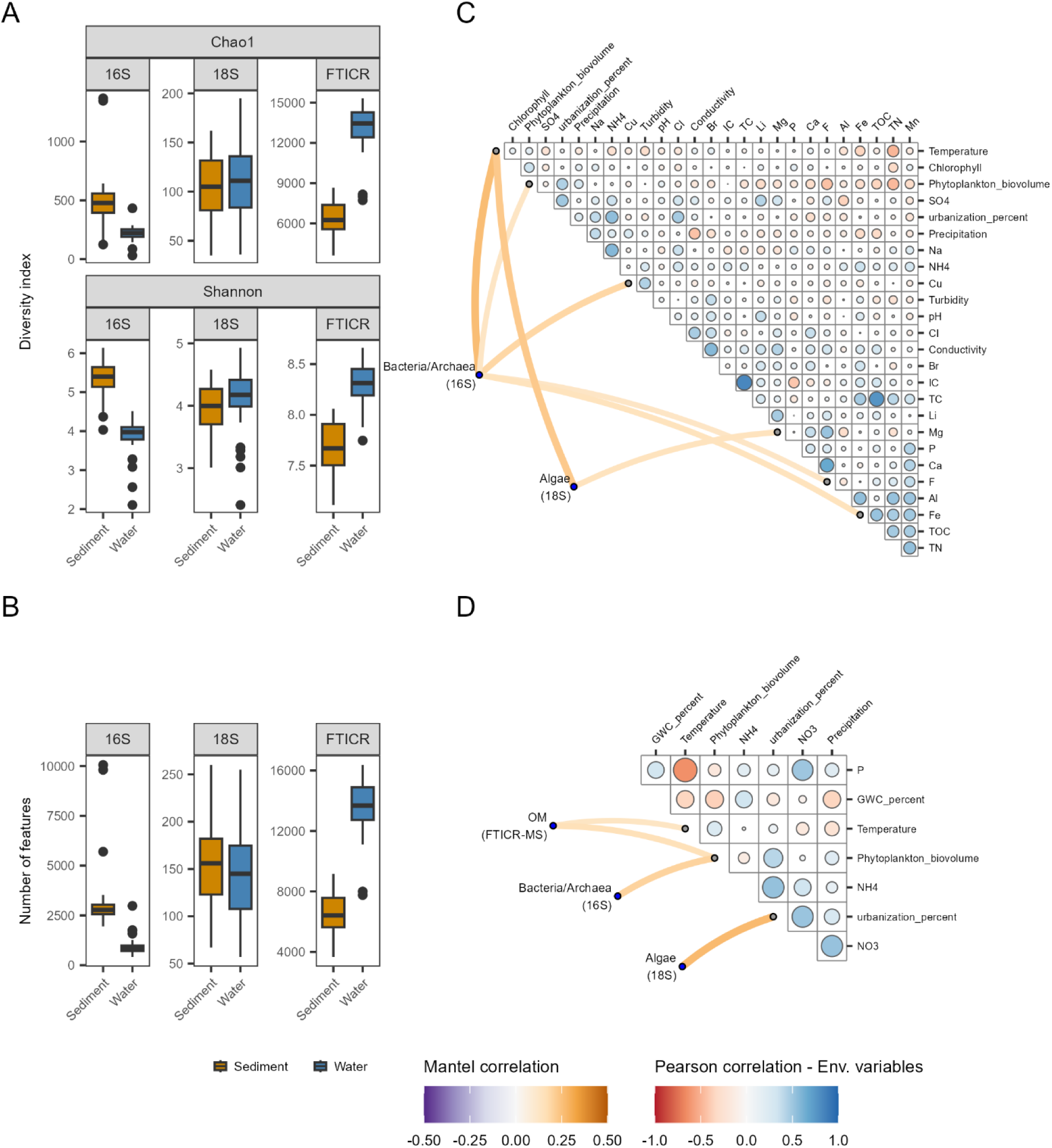
A) Boxplots showing the alpha diversity indexes (Chao1 and Shannon) calculated for the datasets of each sample type. B) Boxplots showing the number of features that were found in each dataset. C) Combined plot of Mantel test and Pearson correlation heatmap for water samples. D) Combined plot of Mantel test and Pearson correlation heatmap for sediment samples. C & D) The circles within the triangular matrix correspond to Pearson correlation strength among environmental variables, while lines illustrate Mantel test results between environmental variables and organic matter profiles, microbial communities and algal community structures. Only significant Mantel results are shown (p-value < 0.05).

**Figure 3.**
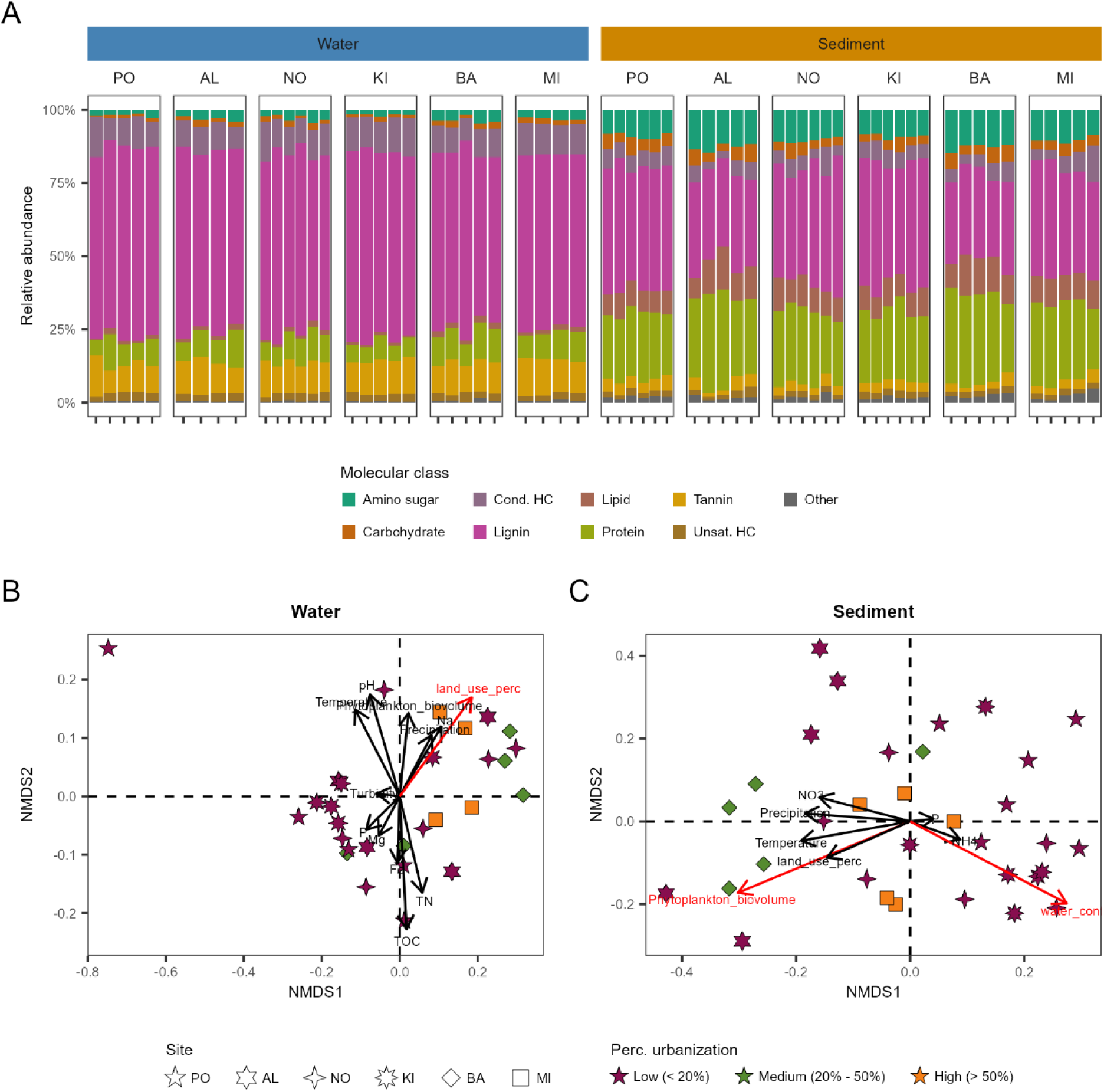
Organic matter profiling. A) Bar plots showing the relative abundance of the putative molecular classes of the detected peaks. B) NMDS ordination of Bray-Curtis distances between the OM profiles of water samples. C) NMDS ordination of Bray-Curtis distances between the OM profiles of sediment samples. Environmental and biogeochemical measurements were fitted to both ordinations and are shown as arrows in the plot. Red arrows indicate a significant fit.

Bray–Curtis beta-diversity revealed that DOM composition responded to different facets of urbanization in benthic vs pelagic environments. In water, the level of urbanization explained 16% of variation in DOM composition (PERMANOVA p-value 0.025), with temperature explaining an additional 7% of variation, albeit with marginal significance (PERMANOVA p-value 0.062) (Figure 3B). Mantel tests showed no significant association between water DOM composition and any biogeochemical measurement. In sediments, DOM composition was significantly explained by urbanization (PERMANOVA: R² = 0.165, p = 0.002), phytoplankton biovolume (R² = 0.083, p = 0.022), and temperature (PERMANOVA R² = 0.095, p = 0.007; Figure 3C). Mantel tests further confirmed this pattern: sediment DOM dissimilarity was significantly associated with phytoplankton biovolume (r = 0.136, p = 0.019) and temperature (r = 0.146, p = 0.018) (Figure 2D). Because temperature and phytoplankton biovolume covaried negatively with TOC, TN, and other micronutrients in the water (e.g., Al, Fe, Mn, Na, NH_4_, SO_4_; Table 1), the apparent temperature effect likely reflects broader seasonal shifts in nutrient availability and associated bloom dynamics rather than direct thermal control of DOM composition.

For bacterial and archaeal communities, 16S rRNA amplicon sequencing recovered 112,230 ASVs across all samples (n = 71), with more ASVs detected in sediments than in the water column. While DOM diversity was similar between the water-column and sediments, sediments had higher bacterial/archaeal diversity (Figure 2A and 2B).

Taxonomically, pelagic communities were dominated by five major classes Actinobacteria, Alphaproteobacteria, Bacteroidia, Cyanobacteriia, and Gammaproteobacteria. Beta-diversity analysis showed that 6% of variation in the pelagic bacterial/archaeal communities was influenced by temperature (PERMANOVA: p = 0.009; Figure 4B). Mantel tests revealed that this temperature signal was accompanied by broad associations with temperature and phytoplankton biovolume, as well as some dissolved ions (Cu: r = 0.140, p = 0.010; Fe: r = 0.143, p = 0.026; F: r = 0.112, p = 0.023; Figure 2C). These multiple correlated variables likely further reflect how broader, seasonally covarying gradients in catchment inputs and ionic composition can modulate pelagic microbial composition.

**Figure 4.**
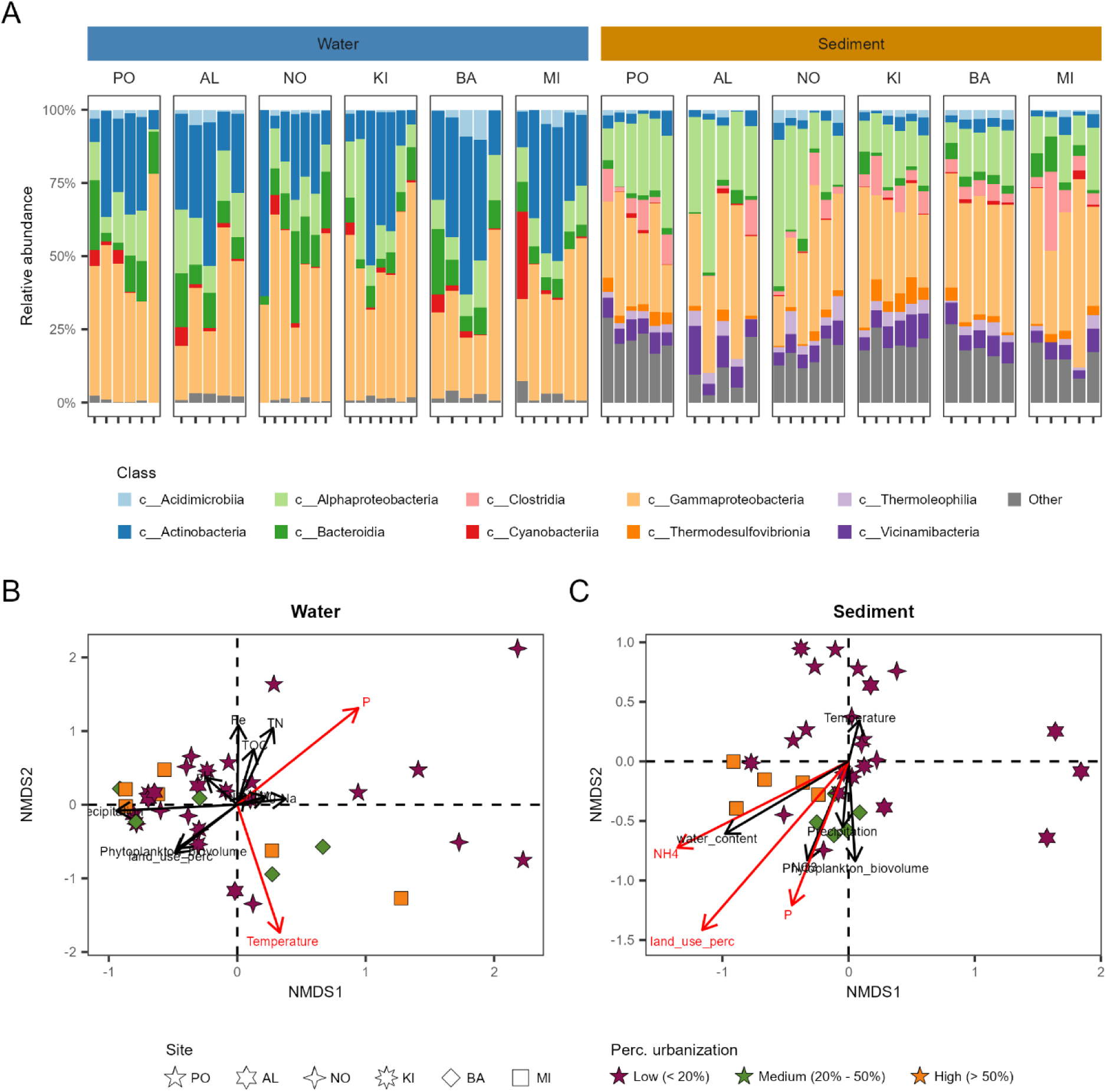
Bacterial and archaeal community profiling. A) Bar plots showing the relative abundance of the top ten most abundant taxonomic classes detected. B) NMDS ordination of Bray-Curtis distances between the bacterial and archaeal profiles of water samples. C) NMDS ordination of Bray-Curtis distances between the bacterial and archaeal profiles of sediment samples. Environmental and biogeochemical measurements were fitted to both ordinations and are shown as arrows in the plot. Red arrows indicate a significant fit.

In contrast, benthic communities were mainly dominated by Gammaproteobacteria and Alphaproteobacteria only, with a substantial fraction of the microbiome (often ∼20–25%) composed of a variety of low-abundance classes in some samples (Figure 4A). Benthic bacteria/archaeal community composition was only significantly associated with urbanization (PERMANOVA: R² = 0.113, p = 0.001; Figure 4C). Mantel tests showed that phytoplankton biovolume was the strongest environmental correlate of sediment bacterial/archaeal dissimilarity (r = 0.296, p = 0.005) (Figure 2D).

For algae, 18S rRNA amplicon sequencing detected 3,090 ASVs across all samples (n = 71). Alpha diversity indices were broadly similar between water and sediments, but community composition was markedly different (Figure 2A, 2B). Water communities were dominated by Chrysophyceae and an unclassified Ochrophyta lineage, with Mediophyceae also abundant at some sites. Conversely, sediments showed higher relative abundances of Chlorophyta (notably Chlorophyceae and Trebouxiphyceae), with Coscinodiscophytina and other diatoms contributing substantially in some samples (Figure 5A). Notably, despite these pronounced compositional differences between compartments, algal community variation was associated with the same potential drivers in both habitats. For pelagic algal communities, temperature (PERMANOVA R² = 0.079, p = 0.001) and urbanization (PERMANOVA R² = 0.088, p = 0.004) were significant predictors of community variation (Figure 5C), with Mantel tests also linking pelagic algal dissimilarity to temperature (r = 0.228, p = 0.005).

**Figure 5.**
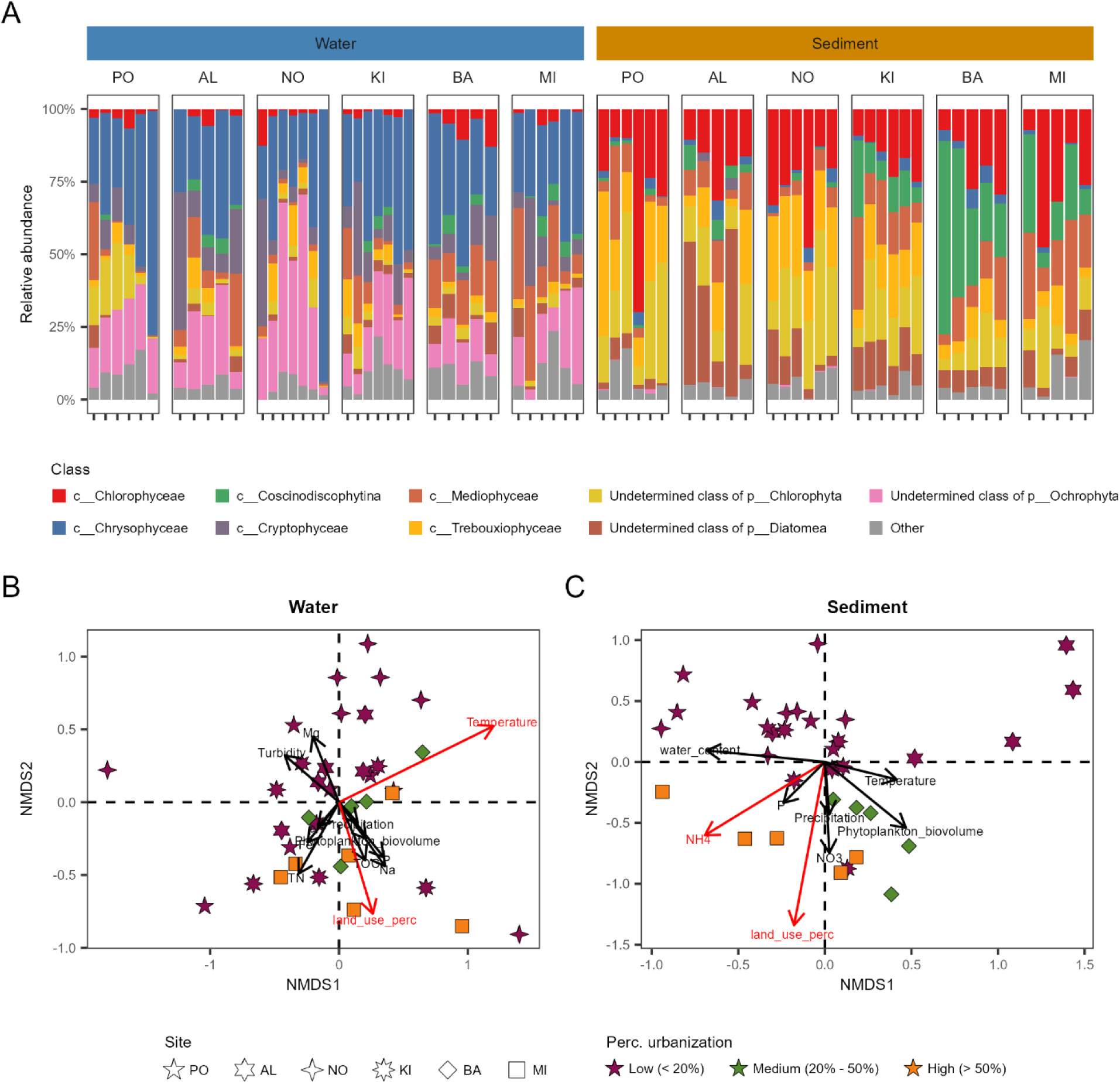
Algal community profiling. A) Bar plots showing the relative abundance of the top ten most abundant taxonomic classes detected. B) NMDS ordination of Bray-Curtis distances between the algal profiles of water samples. C) NMDS ordination of Bray-Curtis distances between the algal profiles of sediment samples. Environmental and biogeochemical measurements were fitted to both ordinations and are shown as arrows in the plot. Red arrows indicate a significant fit.

Benthic algal communities responded to these same factors, with variation similarly linked to urbanization level (PERMANOVA R² = 0.167, p = 0.001) and temperature (PERMANOVA R² = 0.047, p = 0.021). Here, Mantel tests indicated that the urbanization gradient was the strongest correlate of sediment algal dissimilarity (r = 0.269, p = 0.001; Figure 2B). Thus, even though algal community composition differed substantially between the water column and sediments, both compartments appeared to be structured by a shared set of environmental drivers, chiefly temperature and urbanization.

### Bloom intensity reshapes cross-domain microbial–algal–metabolite association networks

Changes in pelagic communities appeared to be largely transient and mediated through seasonally varying processes. For example, phytoplankton biovolume increased initially but decreased by the end of the sampling period (Supplementary Figure 1). Because phytoplankton abundance may be stimulated by urban runoff, as indicated by higher phytoplankton peaks in more urbanized sites (Supplementary Figure 1), we investigated how phytoplankton biovolume itself reshapes the ecological interactions among microbes, algae, and organic matter in the pelagic compartment through the use of SparCC association networks (Figure 6A).

**Figure 6.**
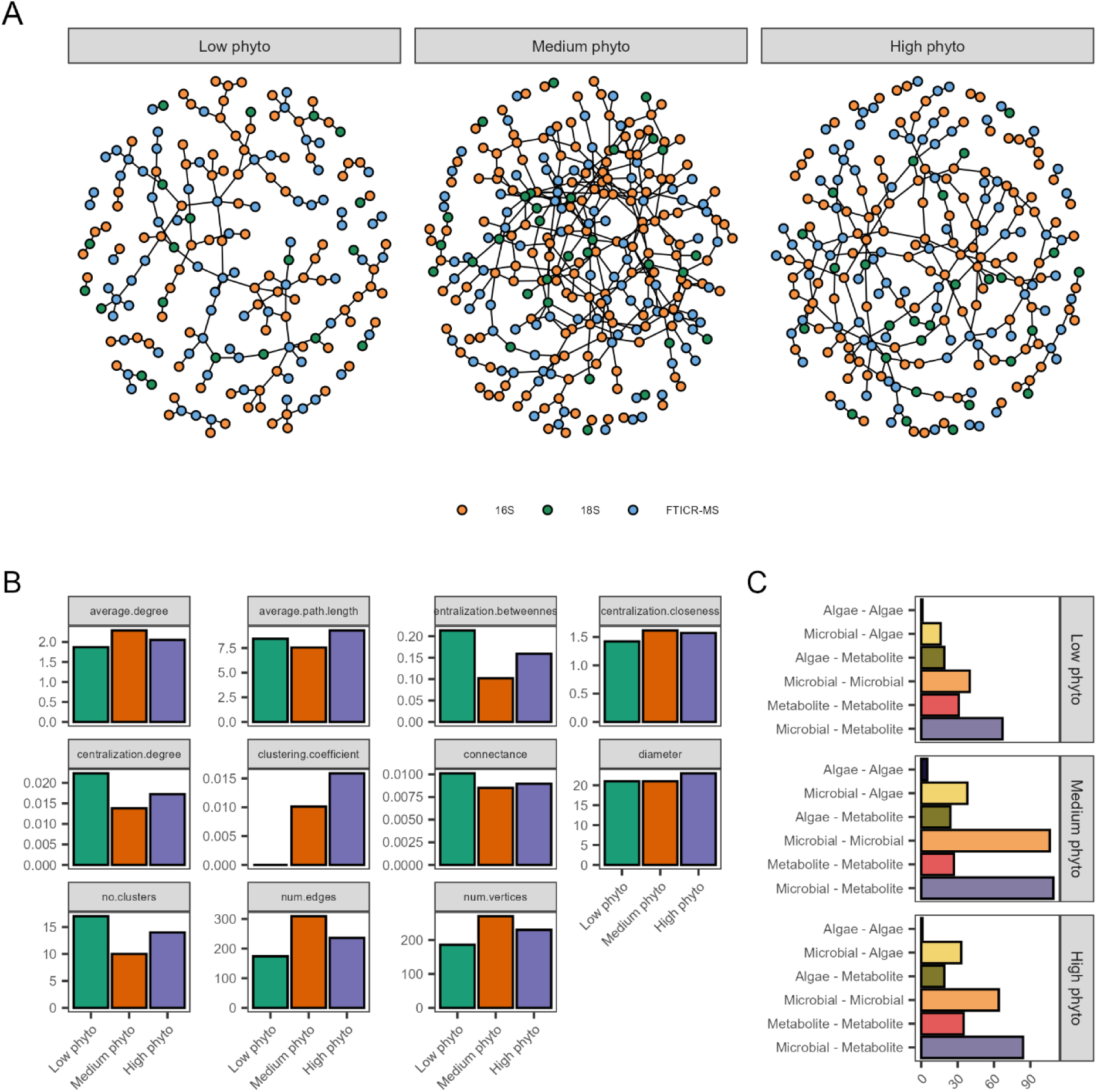
A) SparCC Correlation networks of water samples based on the phytoplankton content. Nodes represent different features; edges represent high Pearson correlations (abs(r > 0.8)) that were significant (P-value < 0.05). B) Statistics of the correlation networks. C) Number of edges between each type of data in the networks

Network size and connectivity differed with phytoplankton biovolume. The coassociation network during medium-sized blooms was the largest and most connected, comprising 270 vertices and 310 edges (average degree = 2.29), compared to the other networks during periods with low or high bloom intensities (Figure 6B). Despite broadly similar connectance values among networks, the increase in vertices and edges from low to medium and high phytoplankton biovolume content indicates a greater number of inferred cross-domain associations under phytoplankton-influenced conditions.

Cross-domain (i.e., bacteria/archaea - algae; bacteria/archaea - metabolite; or algae - metabolite) associations also changed with phytoplankton biovolume. Relative to associations occurring during low phytoplankton biovolume, coassociation networks during medium-sized and large blooms showed pronounced increases in microbial– microbial edges and microbial–metabolite edges (Figure 6C). Even though algal interactions with bacteria/archaea and/or DOM were less common, they nevertheless increased slightly between low and medium- and high-bloom phytoplankton networks. Lastly, metabolite–metabolite associations were observed across all phytoplankton groupings suggesting persistent DOM dynamics despite changes in phytoplankton biovolume.

### Identifying impacts and indicators of phytoplankton biovolume and urbanization

We applied two complementary approaches, Threshold Indicator Taxa Analysis (TITAN2) (Baker et al., 2025) and multiblock sparse partial least square discriminant analysis (sPLS-DA) (Lê Cao et al., 2011), to identify taxa and DOM molecules with consistent directional responses across the phytoplankton biovolume (for the pelagic habitat) and urbanization gradients (for the benthic habitat).

First, we applied TITAN2 to identify microbial taxa and DOM molecules whose changes in abundance may indicate ecological community thresholds along the phytoplankton biovolume gradient in the water column. We revealed a total of 842 DOM molecules, 20 bacterial and 30 algal taxa that showed consistent directional responses in coordination with phytoplankton biovolume changes (Figure 7A). The large number of FTICR-MS indicators suggests that increases in phytoplankton abundances are likely accompanied by shifts in multiple DOM sub-pools. Different types of carbohydrate-, condensed hydrocarbon-, lignin-like compounds were found as indicators of low levels of phytoplankton, with protein-like, tannin-like and other lignin-like compounds increasing at medium levels of phytoplankton biovolume suggesting a distinct chemical transition related with the amount of phytoplankton present in the water column. In the 16S amplicon data, low-phytoplankton conditions were characterized by Gammaproteobacteria, Bacteroidia and Alphaproteobacteria with medium and higher phytoplankton conditions being associated with Actinomycetota-related taxa, specifically Acidimicrobiia and Actinobacteria. This pattern suggests a shift from a Gammaproteobacteria-associated baseline community at low phytoplankton abundance toward Actinomycetota lineages as phytoplankton increase. Finally, for the 18S rRNA gene sequences, taxa like Chryptophyceae were enriched in low phytoplankton conditions while lineages such as Trebouxiophyceae, Chrysophyceae, and Coscinodiscophytina were found to become enriched at medium and high phytoplankton levels.

**Figure 7.**
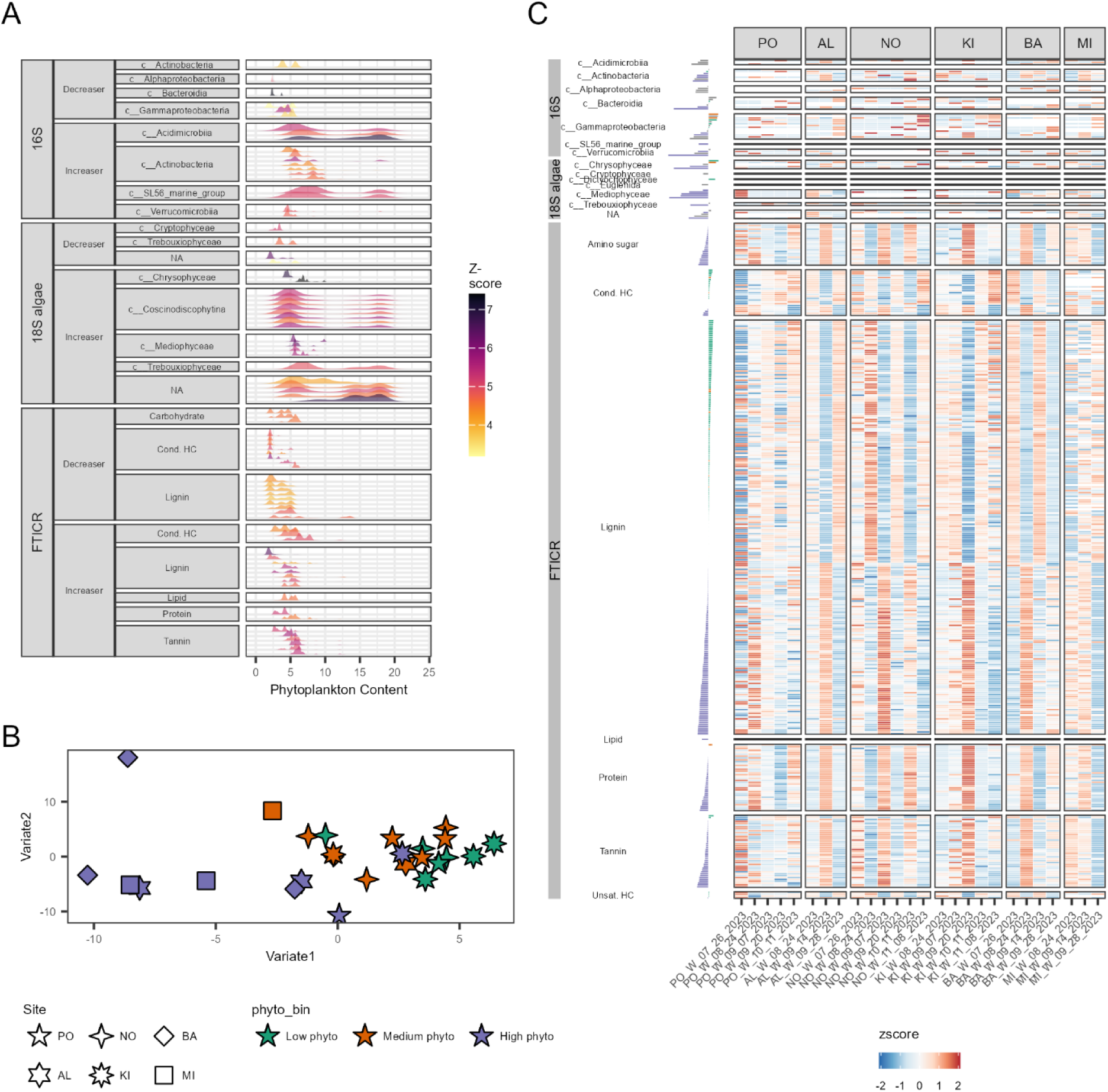
A) Ridge plot of the species identified by TITAN2 as indicators of the change in phytoplankton content. Figure includes all indicators found for the 16S and 18S dataset, but only the top 8 increasers and decreasers of the FTICR-MS dataset. B) Multiblock sPLS-DA ordination plot of water samples using the three datasets 16S, 18S and FTICR-MS. Colors represent the phytoplankton content. C) Heatmap showing the normalized abundances (z-score) of the features found to distinguish between the samples having low, medium and high phytoplankton content. The bars in the Y axis represent their importance across Variate 1.

To complement these results, sPLS-DA was applied to select a reduced set of molecular features (e.g., ASVs or molecular formulas) across the FTICR-MS, 16S, and 18S datasets that allow discrimination between different levels of phytoplankton biovolume (Low, Medium and High), providing actionable biomarkers for monitoring and early warning (Figure 7B). By integrating features from all three datasets, the model defined a multi-omics fingerprint of phytoplankton content, in which the combined signal of DOM molecules, bacterial/archaeal ASVs, and algal taxa jointly discriminated among biovolume levels more robustly than any single dataset alone. The variables composing this fingerprint indicated that changes in phytoplankton biovolume are characterized by coordinated, cross-domain shifts spanning algal taxa, bacterial/archaeal ASVs, and specific DOM molecules (Figure 7C), underscoring that the molecular, microbial, and chemical layers respond in concert. This integrated fingerprint thus offers a concise, mechanistically interpretable signature that could be leveraged for monitoring and early warning of bloom development.

Discriminant features derived from 16S rRNA gene sequences comprised several major bacterioplankton classes typically associated with organic matter cycling during blooms. Taxa from Actinobacteria and Acidimicrobiia, alongside Bacteroidia, Cyanobacteriia, and Alphaproteobacteria were associated with higher phytoplankton content, while taxa from Gammaproteobacteria were mostly associated with lower phytoplankton content. Conversely, almost all of the discriminant ASVs identified from the algal communities were associated with higher phytoplankton content, including representatives from taxa such as Mediophyceae, Chrysophyceae, Chlorophyceae, Trebouxiophyceae, and Cryptophyceae. Similar to the algal markers, most DOM indicators were also highly associated with high phytoplankton biovolume. These molecular formulas included mostly lignin-like molecules, but also several amino sugar-, carbohydrate-, lipid- and tannin-like molecules indicating that phytoplankton-level discrimination was associated with shifts in multiple DOM sub-pools rather than a single compound class. These phytoplankton abundance-associated indicators characterize the transient molecular and microbial signatures of urbanization driven phytoplankton increasing events in the water column.

Next, since our earlier analyses indicated direct effects of urbanization land-use in sediments, we used TITAN2 to characterize the persistent molecular and microbial imprints of urbanization by identifying potential impacts derived from the DOM, bacterial/archaeal, and algal communities. In total, TITAN2 identified 1055 indicator DOM molecules, 43 bacterial/archaeal indicators, and 45 algal indicators. Most DOM indicators were found as markers of low urbanization (Figure 8A). Consistent with this, many lignin- and tannin-like compounds were indicators of low urbanization, whereas some lipid-, amino sugar-, and protein-like molecules were markers of high urbanization. Algal indicators belonged to Chlorophyceae, Coscinodiscophytina, Mediophyceae, and Euglenida were mostly found to become enriched when moving from low to medium urbanization levels, while those from Trebouxiophyceae and other Chlorophyceae were enriched at lower urbanization levels. For the bacterial and archaeal community, Gammaproteobacteria, Nitrospiria and Bacteroidia were consistently associated with increased urbanization. In contrast, Alphaproteobacteria taxa were largely enriched in less urbanized samples.

**Figure 8.**
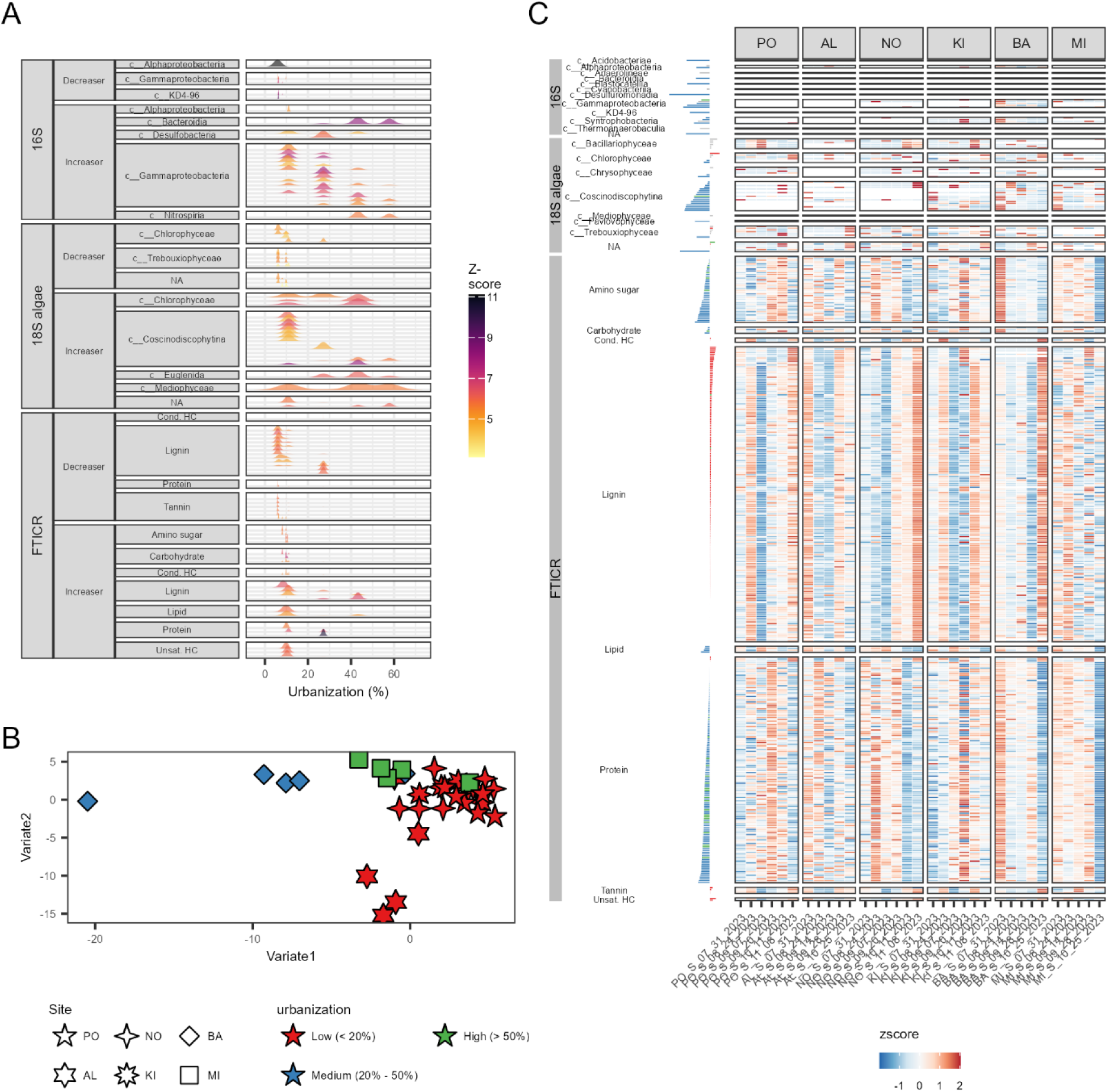
A) Ridge plot of the species identified by TITAN2 as indicators of the change in the level of urbanization. Figure includes the top 8 increasers and decreasers for all the datasets. B) Multiblock sPLS-DA ordination plot of sediment samples using the three datasets 16S, 18S and FTICR-MS. Colors represent the urbanization percentage. C) Heatmap showing the normalized abundances (z-score) of the features found to distinguish between the samples from low, medium and high urbanization environments. The bars in the Y axis represent their importance across Variate 1.

Multiblock sPLS-DA was also applied to determine potential markers of urbanization in sediments to select a reduced set of features across the FTICR-MS, 16S, and 18S datasets that allow discrimination between different levels of urban land use. The resulting discriminant components selected a compact set of variables whose loadings separated low-, medium-, and high-urbanization sites (Figure 8B). Discriminant taxa belonging to the Gammaproteobacteria, Alphaproteobacteria and Bacteroidia were found to be enriched in medium urbanization samples (Figure 8C). In the algal block, discriminant taxa associated with medium and high urbanization included multiple Coscinodiscophytina. Meanwhile, Chlorophyceae taxa were enriched in low urbanization samples. The identified discriminants from the FTICR-MS block further indicated systematic DOM differences across the urbanization gradient. Medium and high-urbanization sediments were associated with a suite of protein-like and amino-sugar-like formula. Low-urbanization sediments, in contrast, were enriched in lignin-like, tannin-like, and unsaturated-hydrocarbon-like compounds.

## Discussion

A central finding of our study is that land use drives temporally decoupled biogeochemical and microbial responses across pelagic and benthic estuarine habitats. Pelagic organic matter and microbial communities are shaped primarily by transient, seasonally forced dynamics, such as temperature shifts, nutrient pulses, and changes in phytoplankton biovolume, which are influenced by urban catchment inputs. In the water column, urbanization appeared to act largely indirectly, with more urbanized sites exhibiting higher phytoplankton peaks (Supplementary Figure 1) that, in turn, reshaped cross-domain microbial–algal–metabolite associations and left coordinated, but transient, molecular and microbial signatures. In contrast, benthic OM and microbial assemblages showed a direct and comparatively stable imprint of urban land use, consistent with persistent and cumulative urban pressures such as chronic nutrient loading, contaminant accumulation, and habitat modification. The enrichment of N-containing compounds (e.g., lipid-, amino sugar-, and protein-like molecules) in more urbanized sediments likely reflects the greater delivery of nitrogen from anthropogenic sources characteristic of urban catchments, such as fertilizer runoff, wastewater inputs, and the deposition of nitrogen oxides from human pollution (Lyu et al., 2021).

Conversely, the more rural sites were characterized by structurally and chemically complex OM (e.g., lignin- and tannin-like compounds) potentially from terrestrial-plant origin, which is generally more resistant to microbial degradation (Hopper et al., 2025; Mueller et al., 2021). This compositional contrast has important biogeochemical implications: the readily bioavailable, N-rich substrates accumulating in urbanized sediments may support a greater diversity of, or shift toward different, microbially mediated biogeochemical processes, including those that cycle and potentially emit nitrogen as greenhouse gases (R. Wang et al., 2025). Together, these habitat-specific responses (i.e., transient, phytoplankton-mediated dynamics in the water column versus persistent, cumulative imprints in the sediments), illustrate how a single urbanization gradient manifests differently across compartments. This persistent-versus-transient duality provides a unifying framework for interpreting the multi-omics patterns observed across all three datasets and both habitats (Figure 9).

**Figure 9.**
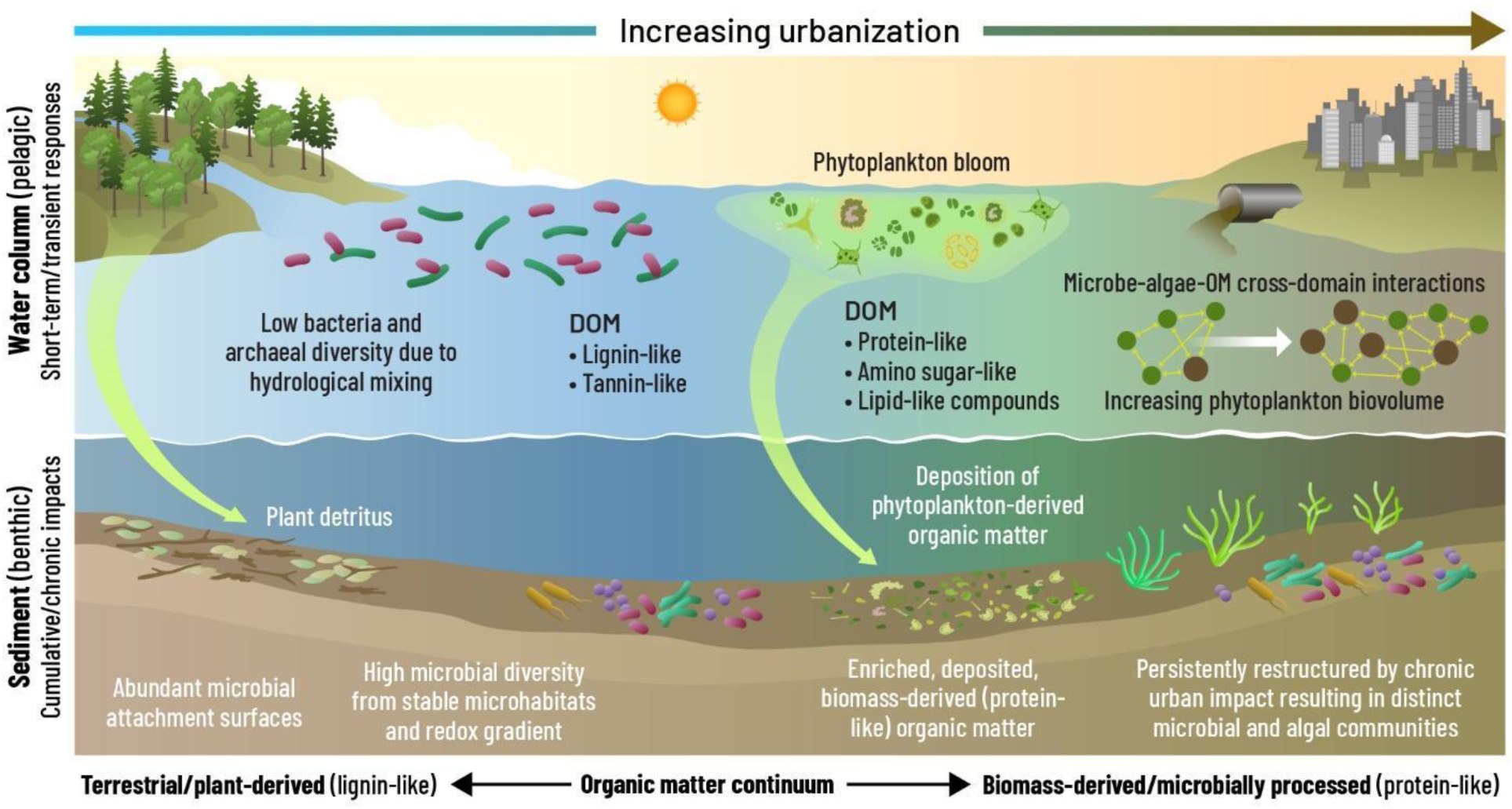
Conceptual model of how urbanization and phytoplankton dynamics reshape dissolved organic matter (DOM) and microbial communities across a freshwater estuary. The model proposes that urbanization and phytoplankton dynamics reshape estuarine ecology through two interacting pathways along an increasing urbanization gradient, from non-urban (left) to urban (right): rapid responses to resource pulses in the pelagic habitat (water column) and slower, persistent, and cumulative effects in the benthic habitat (sediments). In the water column, non-urban conditions support a lignin-and tannin-like DOM pool (most likely due to vegetation inputs), with a lower bacterial and archaeal diversity. Under urban conditions, urban inputs may stimulate phytoplankton blooms result in a more labile biomass-derived and microbially processed DOM pool (containing protein-, amino sugar-, and lipid-like compounds). Throughout the water column, increase in phytoplankton biovolume results in increased and more complex microbe-algae-OM interactions. Downward arrows organic matter deposition towards the sediment, like plant detritus in non-urban conditions as well as phytoplankton derived organic matter across the whole estuary showcasing an example of benthic-pelagic coupling. Chronic urban impacts persistently restructure sediment microbial and algal communities potentially due to sediment organic matter shifts from terrestrial/plant-derived, lignin-like material toward enriched, deposited, biomass-derived, protein-like material alongside the urbanization gradient. Together, the model emphasizes that estuarine responses to urbanization emerge from shifts in the strength and direction of interactions among algae, microorganisms, and organic matter, reflecting transient reorganization in the water column and cumulative, chronic, and integrative change in the sediments.

Overall, our results support that land use impacts on pelagic chemistry fluctuate with the intensity of transient terrestrial inputs (Mallin et al., 2009). The negative correlation of temperature with carbon and nutrient concentrations (TOC and TN), in particular, implies that the observed temperature associations likely represent co-occurring seasonal processes, such as external loading, precipitation, water-column mixing, and nutrient availability (Wu et al., 2014), which are also modulated by urban land use (Valtanen et al., 2014). In the SLRE, hydrological conditions stabilize as river discharge declines from spring into the summer baseflow, promoting passive settling of suspended material, reduced turbidity and carbon loading, longer water residence times, and warmer, more stable water-column conditions. These conditions combined with calm winds, and low-flow regimes documented during the 2023 and 2024 late-summer drought periods (Nicklay et al., 2025), likely resulted in nutrient accumulation and water-column stability. The seasonal dynamics of TN are further shaped by the estuary’s role as a nitrogen removal hotspot, where optimal denitrification conditions, sufficient NO_3_–N, organic-rich sediments, warm temperatures, and long water residence times, creates uniquely high denitrification potential during summer baseflow (Loken et al., 2016). Similarly, the correlations of the bacterial/archaeal community variation to several ions/metals (e.g., Cu, Fe), suggest sensitivity to substrate availability and trace-metal chemistry (J. Zhao et al., 2021) associated with catchment inputs (Jones et al., 2020). However, it is important to note that the most urbanized sites in our study also correspond to lower estuary positions, introducing an estuarine gradient (e.g., differences in ionic composition, water clarity, and hydrodynamics) that may partially confound the urban land-use signal.

Consistent with previous studies (X. Li et al., 2023; Valle et al., 2020; H. Wang et al., 2023), the strongest signal in microbial, algal, and DOM profiles was the separation between pelagic and benthic profiles, reinforcing that these two habitats are governed by different ecological and biogeochemical constraints (Nevers et al., 2020; L. Zhang et al., 2021). Below, we discuss how land use manifests in each compartment, first addressing the indirect, transient influences that structure pelagic composition, and then the direct, persistent mechanisms that shape benthic communities.

The higher chemodiversity of lignin-and tannin-like formulas in the pelagic habitat (Figure 2A, 3A) is consistent with plant terrestrial inputs (Garayburu-Caruso et al., 2020; Hedges & Mann, 1979; Mueller et al., 2021), resulting in a more heterogeneous and continuously renewed DOM pool. The PERMANOVA results further reinforced this interpretation as temperature and urbanization were linked to DOM variation, suggesting that seasonally mediated processes dominate pelagic OM turnover. This is consistent with urban catchments delivering terrestrial inputs in episodic pulses that vary with precipitation, runoff intensity, and seasonal cycles, producing a dynamic chemical environment (Valtanen et al., 2014). In the St. Louis River Estuary (SLRE), where approximately 85% of the landscape remains forested with extensive wetlands, natural land cover likely moderates background nutrient export through retention, infiltration, and reduced runoff (Bomier et al., 2021). As a result, urban inputs become concentrated within small sub-catchments near the lower estuary, increasing the contrast between terrestrially derived DOM and anthropogenic inputs. This landscape configuration likely enhances the detectability of pulsed, seasonally variable urban loadings into the pelagic DOM pool.

In sediments, there were fewer DOM molecules than in the water-column, but a higher proportion of protein-like, lipid-like, and amino sugar-like material (Figure 3A), suggesting accumulation of microbial OM (E. C. Freeman et al., 2024; Mueller et al., 2021). Long-term sedimentary processes (i.e., accumulation, erosion, and deposition) increase nutrient content and moisture (Berhe & Kleber, 2013; L. Zhang et al., 2021), promoting increased microbial biomass and diversity (Du et al., 2020). These processes also make sediments inherently suited to integrate the cumulative effects of urbanization, as contaminants, nutrients, and organic inputs from urban catchments accumulate over time (Kali et al., 2025) with important implications for both monitoring and biogeochemical function. For example, benthic DOM and microbial composition may serve as robust, time-integrated indicators of urban impact that are less sensitive to the transient fluctuations characteristic of the water column, and therefore particularly valuable for detecting chronic, long-term stress. At the same time, the accumulation of bioavailable, N-rich microbial OM in urbanized sediments could sustain elevated rates of microbial metabolism and reconfigure key biogeochemical processes, potentially enhancing nutrient regeneration and the emission of nitrogen-based greenhouse gases from these depositional zones (Murray et al., 2015).

Accordingly, benthic organic matter and microbial composition were most strongly structured by urbanization, particularly for benthic algal communities, where watershed-scale nutrient transport driven by urban land use contributes to persistent restructuring (Hopper et al., 2025). In contrast, microbial communities from the pelagic habitat appeared more responsive to transient dynamics. For example, we observed lower bacterial and archaeal diversity in pelagic communities than in benthic communities (Cole et al., 2013; B.-W. Feng et al., 2009; Ibekwe et al., 2016; H. Wang et al., 2023) may reflect several interacting factors that distinguish these two habitats. For example, sediments provide abundant surfaces for microbial attachment, resulting in denser biomass and higher diversity compared to the pelagic habitat where cells are more sparsely distributed (Graham et al., 2016). Moreover, the physical structure and relative stability of the sediment matrix create a mosaic of microhabitats and redox gradients that promote niche diversification (Graham et al., 2017), compared with the pelagic environment where hydrological mixing can exert a homogenizing effect on the microbial assemblages (Fortunato et al., 2012) reducing local diversity. Together, these dynamics point to more susceptible, less species-rich pelagic bacterial communities turning over rapidly in response to transient urban-derived inputs (Heqing Huang et al., 2021; X. Li et al., 2023; Nelson, 2009). Pelagic algal assemblages showed a similar dual sensitivity: both temperature and urban land use were significant predictors of pelagic algal composition, suggesting that these communities respond to transient seasonal changes and direct urban pressures simultaneously. The additional association with magnesium further reflects ionic changes associated with catchment inputs that influence phytoplankton composition.

The stronger association of benthic communities with urban land use likely reflects inherent sediment properties, including the capacity to adsorb contaminants and serve as long-term nutrient sources (Kalev & Toor, 2018), as well as their role as microbial reservoirs less sensitive to short-term hydrodynamic change (Crump et al., 2012; L. Zhang et al., 2021). Beyond these intrinsic properties, benthic communities were also shaped by inputs from the overlying water column. Specifically, the relationship between sediment bacterial/archaeal dissimilarity and phytoplankton content points to a benthic-pelagic coupling mechanism: urban nutrient inputs stimulate phytoplankton biovolume in the water column (Wurtsbaugh et al., 2019), and the resulting phytoplankton-derived organic matter is deposited into sediments, restructuring benthic microbial assemblages (Xiao et al., 2017). Taken together these observations suggest that the urban biogeochemical signal detected in sediments is attributable to localized catchment-scale inputs rather than diffuse watershed-wide disturbance, providing a clear natural baseline against which urbanization impacts can be resolved.

Since OM chemistry and availability are central drivers of biogeochemical cycles in estuaries (Zhuang & Yang, 2018), identifying specific taxa and compounds that respond to urbanization-linked stressors provides both mechanistic insight and candidate biomarkers for environmental monitoring. To resolve this challenge, we next applied a mixed statistical approach to identify individual molecular and microbial features that characterize compartment-specific responses to land use.

TITAN2 results revealed that relationships of phytoplankton with water-column OM and microbial communities are characterized by non-linear threshold responses, suggesting that increases in phytoplankton biovolume trigger discrete ecological reorganizations. This is consistent with algal blooms resulting sudden regime shifts between clear-water states and algal-dominated states (Scheffer et al., 2009), putatively responding to pulsed nutrient inputs that drive the changes in phytoplankton biomass (Batt et al., 2013; Ortiz et al., 2020). While previous work in the SLRE has focused in thresholds from single physicochemical variables, such as DO, pH or nitrogen levels, as triggers of sudden changes in cyanobacteria abundance (Birschbach et al., 2026), our multi-omics approach aims to understand the biological and chemical dynamics during these transitions simultaneously. By jointly identifying the specific algal taxa, bacterial/archaeal taxa, and DOM molecules that shift in concert across the phytoplankton biovolume gradient, we move beyond documenting that thresholds exist to characterizing which components of the ecosystem respond and how they are coupled. This cross-domain resolution reveals that phtyoplankton-associated thresholds are not confined to a single trophic level or chemical pool, but instead propagate across microbial, algal, and molecular compartments, underscoring the whole-ecosystem reach of phytoplankton growth dynamics. These reorganizations are consistent with feedback between primary production, substrate supply, and microbial processing that lead to stepwise changes in ecosystem structure and function (Eigemann et al., 2023; Landa et al., 2016; Luria et al., 2017). In the context of urbanization, the presence of non-linear transitions suggests that even modest increases in nutrient delivery from urban catchments may push pelagic communities past critical thresholds, triggering disproportionate shifts in OM chemistry and microbial composition.

The large number of DOM molecules exhibiting threshold responses, relative to microbial indicators, further suggests that increases in phytoplankton biovolume are accompanied by broad restructuring of the pelagic DOM pool, potentially reflecting algal exudation, cell lysis, and rapid microbial transformation (Chen et al., 2021; Landa et al., 2016). These indicators may point to stage-specific DOM dynamics across bloom intensification. Consistent with our finding that phytoplankton-level discrimination was associated with shifts across multiple DOM sub-pools rather than a single compound class, at higher phytoplankton biovolume algal exudates would accumulate as labile lipid- or protein-like compounds that can be subsequently degraded into recalcitrant lignin- or tannin-like molecules (Jiao et al., 2010; Landa et al., 2016; Q. Zheng et al., 2019). Such indicators would be particularly valuable for detecting transient urban impacts in the water column, where fluctuating phytoplankton abundances make them difficult to capture with conventional monitoring.

Pelagic microorganisms exhibiting threshold responses were consistent with contrasts between baseline microbial communities and the communities that co-occur with high levels of phytoplankton (Mühlenbruch et al., 2018). An increase in phytoplankton biovolume appeared to be associated with the enrichment of Actinobacteriota taxa, that may specialize in the uptake of low-molecular-weight compounds derived from algal exudates (Newton et al., 2011), as compared to low phytoplankton conditions that were associated with many Gammaproteobacteria. For algae, the decline of Cryptophyceae at the low end of the phytoplankton gradient is consistent with their mixotrophic ecology (Unrein et al., 2007), while Trebouxiophyceae and Coscinodiscophytina exhibit metabolic flexibility that allows them to thrive under high-nutrient conditions (Armbrust, 2009; Xiong et al., 2025). Notably, common cryptophytes in the SLRE, including *Cryptomonas* and *Rhodomonas*, are facultative mixotrophs capable of supplementing photosynthesis through phagotrophy (Ballen-Segura et al., 2017), potentially linking bloom-associated shifts in phytoplankton biomass with bacterial turnover and organic matter processing. The threshold behavior of these taxa implies that they could serve as potential biological indicators of changes of phytoplankton biovolume, as their enrichment or decline marks the crossing of critical resource-supply thresholds; consequently, monitoring these indicator taxa may provide early warning of the discrete ecological reorganizations that accompany increases in phytoplankton biovolume.

In contrast to transient pelagic signatures driven by fluctuating phytoplankton abundances, benthic OM and communities had temporally consistent threshold responses to urbanization, regardless of changes in phytoplankton biovolume. The largest number of threshold-like responses occurred during the transition from low to medium levels of urbanization (Figure 8A). This suggests that even small amounts of urban development (around 10% of the surrounding area) may push estuary sediments into a different biogeochemical regime (H. Wang et al., 2023; S.-Y. Wang et al., 2011).The sensitivity of sediments to urbanization contrasts with the water column, where phytoplankton-driven changes were found to be episodic and reversible; in sediments, once urbanization pushes the system past a biogeochemical threshold, the resulting shifts may be difficult to reverse. This underscores that early sediment monitoring is essential to prevent changes in estuarine function caused by urbanization.

While we note that only one site in this study was classified as highly urbanized, and that impacts from a nearby wastewater treatment facility may have been lower in 2023 than other years due to drought (Birschbach et al., 2026), the results presented here still underline that low levels of urbanization have the potential to induce long-term changes in estuarine function via their impacts on sediment habitats.

The molecular indicators found in this study suggest that land use-driven shifts in OM bioavailability, from aromatic, plant-derived compounds toward more labile, N-rich substrates, are accompanied by predictable changes in microbial community composition, linking land-use intensity to the biogeochemical template that governs carbon and nutrient cycling in estuarine sediments. The association of lignin- and tannin-like compounds with low levels of urbanization, is consistent with a stronger contribution of plant-derived, aromatic carbon pools in less urbanized catchments (Hopper et al., 2025; Mueller et al., 2021). In contrast, lipid-, amino sugar-, and protein-like compounds characterized higher urbanization, consistent with greater anthropogenic OM inputs (Lyu et al., 2021). These molecular shifts were mirrored by the microbial community, as microbial indicators such as Gammaproteobacteria, Nitrospiria, and Bacteroidia were found to be associated with increased urbanization, whereas Alphaproteobacteria taxa were characteristic of low-urban sites, which is consistent with the metabolic capabilities of these taxa. Gammaproteobacteria and Bacteroidia are metabolically diverse groups that include many organisms capable of using alternative electron acceptors (e.g., sulfur, iron) (Dong et al., 2021; Z. Zhou et al., 2022) and of degrading organic contaminants (André et al., 2026), traits well suited to the N-rich, contaminant-laden sediments of urbanized catchments. Meanwhile the enrichment of Nitrospiria, key members during nitrogen cycling (Kop et al., 2025), further reflects the elevated nitrogen loading characteristic of urban land use. Conversely, many Alphaproteobacteria are adapted to lower-nutrient conditions and complex, plant- or soil-derived dissolved organic matter (C. Zhou et al., 2026), like those that predominate at less urbanized sites, consistent with their enrichment there.

As the composition and availability of the DOM pool will determine the activity of heterotrophic microbial communities, nutrient regeneration, carbon processing, and energy flow through the ecosystem (Traving et al., 2017; Williams et al., 2010), coordinated shifts across biological and chemical pools carry the greatest consequences for ecosystem function. Thus, we characterized these cross-domain changes, using SparCC association networks and multiblock sPLS-DA.

The systematic changes in network topology provided a systems-level signature of how phytoplankton abundance reorganizes the interactions between algae, bacteria/archaea, and OM in the water column. Increased phytoplankton biovolume, which promotes the heterogeneity of available substrates through exudates and lysates (Chen et al., 2021; Landa et al., 2016), resulted in stronger associations within bacterial and archaeal communities and increased interactions between microbial taxa and DOM molecules. These dynamics are likely compounded by climate variability: in the SLRE, temperature was the most important predictor of cyanobacterial biovolume (Birschbach et al., 2026), and warming trends may increasingly synchronize with land-use-derived nutrient subsidies to intensify phytoplankton-driven network reorganization. Notably, the cross-domain network characterizing periods of moderate phytoplankton biovolume was the most connected and least fragmented, suggesting that greater cross-domain integration may occur at intermediate levels of disturbance (Hou et al., 2024), in a pattern resembling the intermediate disturbance hypothesis (Connell, 1978). In contrast, the lower connectivity and higher fragmentation in the low-phytoplankton network are consistent with weakly linked microbial dynamics under substrate-limited conditions (Hou et al., 2024). Even though algae interactions were rare (Figure 6C), their increase in the medium- and high-phytoplankton networks supports cross-domain coupling between algae, bacteria and OM increasing alongside phytoplankton biovolume (Focardi et al., 2025).

In parallel, we extracted a compact, cross-domain feature set characterizing phytoplankton abundance-driven biogeochemical shifts in pelagic habitats using multiblock sPLS-DA. This approach overcomes limitations such as stochastic nutrient loading, and spatial complexity, which may affect other statistical approaches (Carpenter & Brock, 2006). Many Actinobacteria, Acidimicrobiia, Bacteroidia, Alphaproteobacteria and Cyanobacteriia were associated with higher phytoplankton content, while Gammaproteobacteria were associated with low phytoplankton content, reinforcing the observed transition from a Gammaproteobacteria-associated baseline toward Actinomycetota-enriched communities as phytoplankton biovolume increase. For algae, the most discriminant taxa (e.g., ASVs from Mediophyceae and Chlorophyceae) were associated with higher phytoplankton biovolume, as these fast-growing organisms can exploit nutrient-rich conditions (Lee et al., 2025). Organic matter discriminants associated with high phytoplankton biovolume included mainly lignin-like molecules, but also several amino sugar-, carbohydrate-, lipid- and tannin-like molecules indicating that changes in phytoplankton biovolume was associated with shifts in multiple DOM sub-pools rather than a single compound class. The concordance between both approaches strengthens our confidence in the proposed indicators: TITAN2 identified taxa/compounds that respond consistently along the phytoplankton gradient and highlights the specific thresholds at which these reorganizations occur, while multiblock sPLS-DA demonstrates that a compact subset of discriminant features can classify samples based on their phytoplankton biovolume. Together, these threshold-specific indicators could provide early-warning signals of phytoplankton-driven community reorganization before the system crosses a critical point (Dakos et al., 2012).

Finally, we applied the same multi-omics integration approach to characterize how land-use effects reshape the biological and chemical landscape of estuarine sediments. The multiblock sPLS-DA showed that urban impacts on sediments can be represented by a small set of microorganisms, compounds, and algae. These included several members of Gammaproteobacteria alongside Alphaproteobacteria and Bacteroidia, groups abundant in urban-affected aquatic systems (W. Feng et al., 2022; Gonzalez Mateu et al., 2019) where they play a role in organic pollutant degradation (W. Feng et al., 2022). Among the algal block, lineages such as Coscinodiscophytina, and Mediophyceae, known to thrive in urban estuaries (Ajani et al., 2023), were associated with high urbanization levels. Moreover, higher urbanization sediments were associated with protein-, and amino-sugar–like formulas signaling microbially derived and bioavailable DOM fractions that increase under urban loading (Hosen et al., 2014). Meanwhile low-urbanization sediments were enriched in lignin-/tannin- and condensed-hydrocarbon-like compounds indicating vegetation-derived OM (Lu et al., 2015). Taken together, the multi-omics discriminants reveal that urbanization shifts sediments toward more N-rich, biomass-associated OM pools (Lyu et al., 2021; Wakida & Lerner, 2005) and microbial assemblages capable of degrading urban pollutants (W. Feng et al., 2022), while less urbanized sites retain a stronger aromatic/plant-derived OM signature (Mueller et al., 2021) and distinct microbial/algal communities.

These networks illustrate how nutrient-enriched urban runoff stimulates phytoplankton growth (Y. Zheng et al., 2024), with the resulting organic matter inputs cascading through microbial and chemical networks. The temporary amplification of microbial– metabolite associations likely reflect intensified heterotrophic processing of phytoplankton-derived substrates (Eigemann et al., 2023), accelerating carbon turnover and nutrient regeneration in the water column during these episodic events.

Our conceptual model proposes that urbanization and phytoplankton dynamics reorganize estuarine ecology through two interacting pathways: rapid pelagic responses to resource pulses and slower, cumulative changes in benthic habitat. Consistent with our results showing that pelagic OM and microbial composition were most strongly associated with temperature and phytoplankton biovolume rather than land-use directly, the pelagic environment seems to function as a dynamic reaction space in which runoff, mixing, temperature, and phytoplankton production alter the quantity and composition of available organic substrates over relatively short timescales (Liu et al., 2024; Mannino & Harvey, 2000). These changes restructure microbial communities and their associations with DOM, as evidenced by the increase in cross-domain (microbe-algae-OM) interactions with phytoplankton biovolume. This pattern indicates tighter microbial-DOM coupling under bloom conditions, supporting the view that phytoplankton act as ecosystem engineers by supplying chemically diverse organic matter through exudation and mortality (Kieft et al., 2021).

Our results further suggest threshold-like responses to phytoplankton production. Under low-phytoplankton conditions limited labile carbon likely constrains microbial activity, resulting in smaller, weakly connected networks dominated by a Gammaproteobacteria-associated community. As phytoplankton abundance increases, algal-derived carbon and nutrients expand the number of available metabolic niches, strengthen microbial interactions, and promote closer coupling between microbial taxa and organic matter transformations. Mixotrophic cryptophytes may further enhance these linkages by transferring carbon and nutrients across algal, bacterial, and archaeal communities

Sediments show the ecological memory of these pelagic and watershed processes, integrating repeated inputs of terrestrial material, urban-derived OM, microbial biomass, and phytoplankton detritus (Y. Yu et al., 2012). The enrichment of protein-, lipid-, and amino sugar-like compounds suggests accumulation of microbially processed OM that supports diverse microbial communities and persistent resource gradients. Along with the urbanization gradient, sediment OM shifted from vegetation-derived material toward nitrogen-rich, bioavailable substrates, accompanied by enrichment of Gammaproteobacteria, Bacteroidia, and Nitrospiria. These changes likely favor microbial strategies associated with rapid resource use, nitrogen cycling, and degradation of anthropogenic compounds (H. Li et al., 2025).

Together, these findings highlight strong pelagic-benthic coupling across the estuary. Phytoplankton production supplies organic matter to sediments, whereas urbanization modifies the quality and persistence of substrates in both habitats. Consequently, the water column reflects short-term ecological dynamics, whereas sediments preserve the cumulative effects of land use and repeated phytoplankton deposition. Urbanization therefore reorganizes estuarine ecosystems primarily by altering interactions among phytoplankton, microorganisms, and organic matter rather than by independently affecting each component.

These mechanisms are summarized in Figure 9, which illustrates contrasting non-urban and urban conditions across the pelagic and benthic habitats. In the water column, non-urban conditions are characterized by a highly diverse, lignin-dominated DOM pool and a sparse, homogeneously mixed microbial assemblage of lower bacterial and archaeal diversity, while urban conditions show a more labile (protein-, amino sugar-, and lipid-like) DOM pool. Moreover, promotion of phytoplankton content by urban inputs seems to intensify microbe-algae-OM cross-domain interactions. Deposition of phytoplankton-derived organic matter links the two habitats across the urbanization gradient reinforcing the idea of benthic–pelagic coupling.

Together, our findings demonstrate that urbanization reshapes freshwater estuaries by modifying ecological coupling among phytoplankton, microorganisms, and organic matter across pelagic and benthic habitats. The water column responds rapidly to hydrologic variability and phytoplankton production, whereas sediments integrate longer-term effects of watershed land use and organic matter deposition. The nonlinear responses observed along urbanization and phytoplankton gradients suggest that ecological reorganization may accelerate once environmental thresholds are exceeded. By integrating algae, microorganisms, and organic matter, this study provides a framework for understanding how urbanization modifies carbon and nutrient cycling in freshwater estuaries and for predicting ecosystem responses to continued watershed development, altered hydrology, and changing bloom dynamics.

## Data Availability

The data that support the findings of this study are openly available in Figshare at https://figshare.com/s/ee58704c2354fd3be3cc.

The code for analyzing the data and generating all of the figures can be found in the GitHub repository: https://github.com/Coayala/lake_superior_urbanization

## Acknowledgements

This study was supported by the U.S. Department of Energy, Office of Science, Biological and Environmental Research program Early Career award to EBG. The work was performed by Pacific Northwest National Laboratory, operated by Battelle Memorial Institute for the U.S. Department of Energy under Contract DE-AC05-76RL01830. We thank the Lake Superior National Estuarine Research Reserve, part of the NOAA National Estuarine Research Reserve System (NERRS), for support of this research.

The National Estuarine Research Reserve System is established under Section 315 of the Coastal Zone Management Act, as amended.

## Tables

Table 1. Pairwise Pearson correlation results among physicochemical, hydrological, land-use and biological variables measured in the water dataset. For each variable pair, the table reports the correlation coefficient (r), the associated t-statistic, the p-value, the lower and upper bounds of the 95 % confidence interval, the correlation method used (Pearson) and the source dataset (“water”). Variables include water temperature, chlorophyll a, conductivity, pH, inorganic carbon (IC), total organic carbon (TOC), total carbon (TC), total nitrogen (TN), dissolved ions and trace elements (Al, Br, Ca, Cl, Cu, F, Fe, Li, Mg, Mn, Na, P, NH₄, SO₄), turbidity, catchment urbanization (%), precipitation and phytoplankton biovolume.

